# Prophage-dependent neighbor predation fosters horizontal gene transfer by natural transformation

**DOI:** 10.1101/2020.09.19.304527

**Authors:** Roberto C. Molina-Quiroz, Triana N. Dalia, Andrew Camilli, Ankur B. Dalia, Cecilia A. Silva-Valenzuela

**Affiliations:** Centro de Estudios Científicos, Valdivia, Los Rios, Chile; Department of Biology, Indiana University, Bloomington, IN, USA; Department of Molecular Biology and Microbiology, Tufts University, School of Medicine, Boston, MA, USA

## Abstract

Natural transformation is a broadly conserved mechanism of horizontal gene transfer (HGT) in bacteria (1) that can shape their evolution through the acquisition of genes that promote virulence, antibiotic resistance, and other traits (2). Recent work has established that neighbor predation via Type VI secretion systems (3), bacteriocins (4) and virulent phages (5), play an important role in promoting HGT. Here, we demonstrate that in chitin estuary microcosms, *Vibrio cholerae* K139 lysogens exhibit prophage-dependent neighbor predation of non-lysogens to enhance HGT. Through predation of non-lysogens, K139 lysogens also have a fitness advantage in these microcosm conditions. The ecological strategy revealed by our work provides a better understanding of the evolutionary mechanisms used by bacteria to adapt in their natural setting and contributes to our understanding of the selective pressures that may drive prophage maintenance in bacterial genomes.

**IMPORTANCE:** Prophages are nearly ubiquitous in bacterial species. These integrated phage elements have previously been implicated in horizontal gene transfer (HGT) largely through their ability to carry out transduction (generalized or specialized). Here, we show that prophage-encoded viral particles promote neighbor predation leading to enhanced HGT by natural transformation in the water-borne pathogen *Vibrio cholerae*. Our findings contribute to a comprehensive understanding of the dynamic forces involved in prophage maintenance which ultimately drive the evolution of naturally competent bacteria in their natural environment.

## TEXT

Several bacterial species have evolved to capture DNA (natural competence) as a source of nutrients (6) or to incorporate into their genome to speed their evolution via a process termed natural transformation (NT). *V. cholerae* is a genetically tractable and well-established model organism to study NT. This human pathogen is usually found in association with the chitinous carapaces of zooplankton in estuary and ocean waters (7). *V. cholerae* can utilize chitin as a major carbon and nitrogen source and additionally, this polymer is a required signal for induction of NT (8). In estuarine chitin microcosms, it has been shown that this pathogen can take up multiple large DNA fragments when the exogenous DNA concentration is high (3, 9). Much evidence points to NT and HGT having contributed to the evolution of *V. cholerae* (10).

About 50% of *V. cholerae* clinical isolates carry the temperate kappa phage K139 (11). However, the role of K139 in the ecology of *V. cholerae* has been ill-defined. Here, we explore whether the lytic replication of K139 affects the physiology of *V. cholerae* in chitin microcosms, which mimics the aquatic reservoir for this facultative pathogen.

Using the K139 lysogen E7946, an O1 El Tor *V. cholerae* strain, we first evaluated bacterial replication and production of viral particles over time in chitin microcosms. *V. cholerae* growth was slow and bacterial numbers increased 100-fold by 24h and 1000-fold by 48h of incubation, while K139 plaque forming units (PFUs) increased 3-logs by 6 hours, which is when bacteria were entering exponential growth (**Figure 1A**). Interestingly, we found that insoluble chitin specifically increased K139 PFUs when compared to other carbon sources (**Fig. S1**). Based on these results, we hypothesized that K139 might have a role in the ecology of *V. cholerae* in its environmental reservoir by increasing the competitive fitness of lysogenic strains. To evaluate if K139 is able to kill non-lysogenic strains in co-cultures, E7946 (lysogen) and an isogenic E7946 ΔK139 mutant (non-lysogen) were competed in a chitin estuary microcosm. After 24 h of incubation, the E7946 ΔK139 non-lysogen was outcompeted 8-fold by the E7946 lysogenic strain (**Figure 1B**). To further test if K139 could help lysogens outcompete non-lysogens in this environment, we competed E7946 with a panel of diverse clinical isolates that naturally lack K139. E7946 was able to outcompete all of these clinical isolates (**Figure 1B**). This effect was dependent on K139 viral production, because a E7946 ΔK139 strain was unable to outcompete non-lysogens (**Figure 1B**). These results strongly suggest that K139 plays an ecological role by providing a competitive advantage for lysogenic strains in mixed populations containing non-lysogens. By excising in a small fraction of lysogenic cells, and efficiently killing the neighboring non-lysogenic cells via lytic replication, K139 may allow the remaining lysogenic population to successfully compete for resources in a nutrient-limited estuary environment.

**Figure 1.**
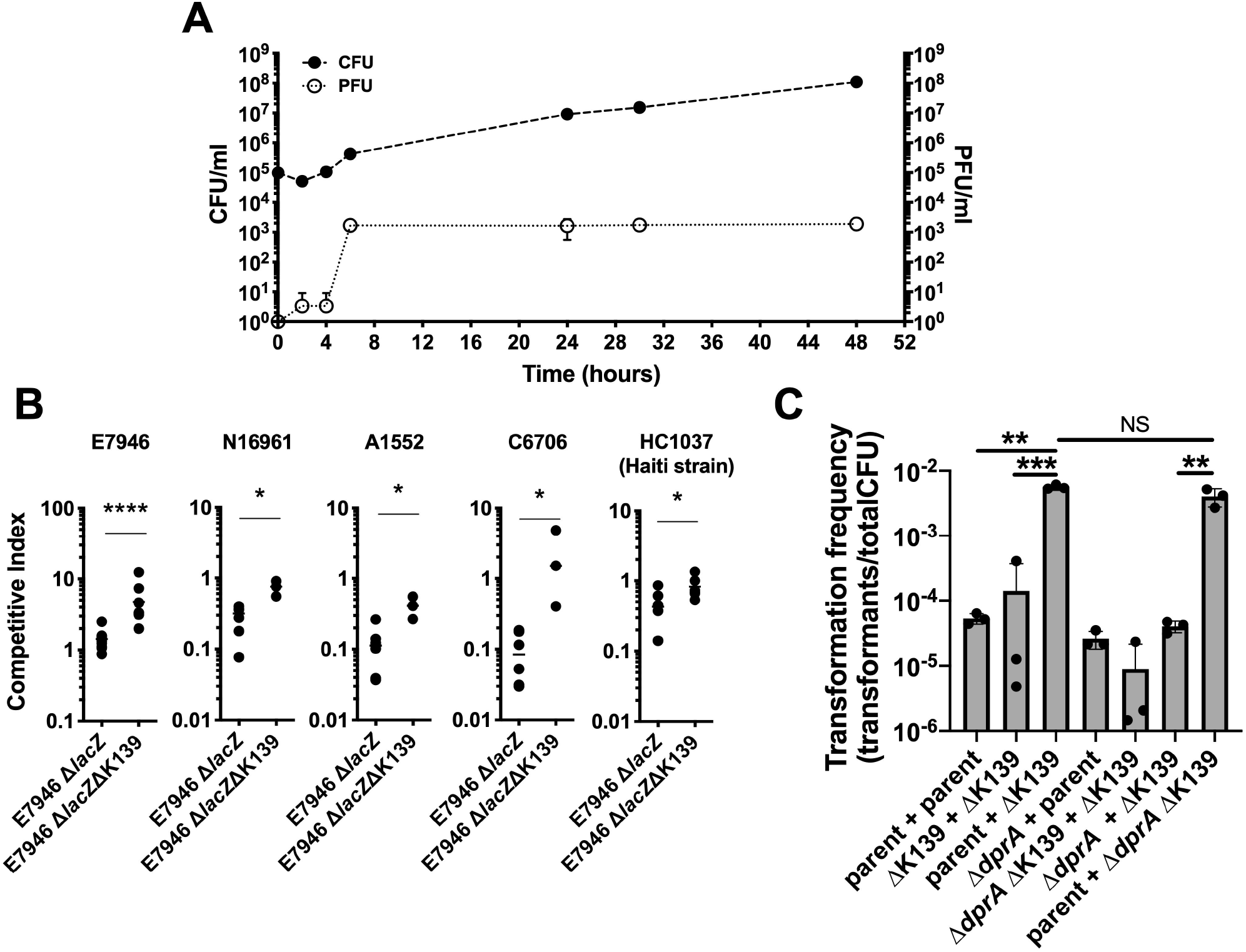
K139 promotes neighbor predation and enhances HGT by NT in chitin microcosms. **A**, *V. cholerae* E7946 growth (CFU, full circles) and K139 phage titer in culture supernatants (PFU, empty circles) were measured in chitin microcosms for 48 hrs. **B**, Competition between *V. cholerae* strains mixed 1:1 was assessed following 24 hrs of growth in chitin microcosms. The strain indicated at the top of each graph was competed with each strain indicated on the X-axis. The competitive index is reported as the ratio of (top strain) / (X-axis strain) in the output divided by the same ratio in the input. Data are from 4 independent experiments and the line within samples denotes the median. The dotted line indicates a CI of 1, which is the value expected if strains compete equally. Statistical comparisons were made by Mann-Whitney test (*p<0.05, ****p<0.0001). **C**, The indicated variants of *V. cholerae* E7946 were co-cultured in chitin microcosms at 30° C for 48 h to assess HGT, which is reported as the transformation frequency (see Methods for details). Data represents three independent biological replicates shown as the mean ± SD. Statistical comparisons were made by one-way ANOVA with Tukey’s post-test on the log-transformed data (**p<0.01, ***p<0.001).

It has previously been shown that some phages can lyse bacterial cells and release intact DNA (5), while other phages degrade host DNA following lysis (12). Release of intact DNA could aid in promoting horizontal gene transfer by NT (5, 13). Here, we have found that K139 promotes neighbor predation in the same chitin microcosm conditions that are required to induce NT. Therefore, we next wanted to test if K139-dependent neighbor predation could promote HGT in a chitin microcosm. To that end, we co-cultured a mixture of strains where each carried an unlinked selection marker at a neutral site in the genome (9). The generation of strains with both selection markers indicated HGT. After 48 h of growth in chitin estuary microcosms, the co-culture containing E7946 (lysogen, wt) and E7946 ΔK139 (non-lysogen), where neighbor predation is expected to occur, showed the highest number of transformants, which was ~100-fold higher than a co-culture of two K139 lysogens or two non-lysogens where neighbor predation is not expected to occur (**Figure 1C**). These results are consistent with neighbor predation promoting HGT.

Prophages have mainly been linked to HGT by transduction (13). However, in chitin microcosms, *V. cholerae* can also undergo HGT via NT. Thus, next we designed a strategy to determine if the HGT observed is attributed to phage transduction or NT. To distinguish between these, we inactivated *dprA* in our *V. cholerae* strains, a gene that is essential for NT but is not required for transduction (14). After 48 h of growth in chitin estuary microcosms, the co-culture containing a NT+ lysogen (E7946) and a NT-nonlysogen (E7946 ΔK139 Δ*dprA*) showed elevated rates of HGT similar to the co-culture containing both NT+ strains (**Figure 1C**). In contrast, a mixture containing an NT-lysogen and an NT+ nonlysogen showed the basal levels of HGT seen in co-cultures where no phage predation occurs (**Figure 1C**). This suggests that the HGT observed is due to the transfer of DNA from the non-lysogen to the lysogen. This is the opposite of what would be expected for phage transduction where a prophage excised from a lysogenic strain would transduce DNA to the non-lysogen. Together, our results point to an adaptative strategy used by lysogenic strains to induce prophage-dependent neighbor predation in order to thrive and also to capture released DNA for HGT in its aquatic environment.

To the best of our knowledge, our work is the first to establish an ecological role for K139 in enhancing *V. cholerae* fitness. We show that K139 can enhance survival of *V. cholerae* lysogens in chitin estuarine environments by neighbor predation of non-lysogens and by driving evolution via HGT. More broadly, our results suggest a novel mechanism by which prophages benefit their lysogenized hosts, which may contribute to the maintenance of these genetic elements in bacterial genomes.

## METHODS

### Bacterial strains and growth conditions

See **Table S1** for a list of all *V. cholerae* strains used in this study. All strains were routinely grown in Luria-Bertani Miller (LB) Broth and agar at 30°C. Where necessary, media was supplemented with erythromycin (10 μg/mL) or trimethoprim (10 μg/mL).

For chitin utilization experiments, cells were grown overnight in LB at 30°C shaking. The next morning, cultures were washed and diluted in 0.7% Instant Ocean (Aquarium Systems) or M9 minimal media. 10^5^ CFU were inoculated in a 1% shrimp shell chitin (Sigma) suspension in 0.7% Instant Ocean (chitin microcosm) and were incubated up to 48h at 30°C statically in 14 ml glass test tubes (Fisher Scientific). Lactate or glucose were added to a final concentration of 0.2% and 0.5% when required. GlcNAc sugars were added at 2mM for pentasaccharides, 3.33 mM for trisaccharides, 5 mM for disaccharides, and 10 mM for monosaccharides. CFU counts were evaluated by serially diluting and plating on LB agar plates.

### Bacteriophage assays

To assay phage titers, supernatants from bacterial cultures were filtered using 0.2 μm filters (Costar). Filtered supernatants were serially diluted and tittered using E7946 ΔK139 (nonlysogenic, susceptible strain) as described in (15). Plates were incubated overnight at 37°C and turbid plaques were counted the next morning.

### Competition and HGT assays

Competition assays were conducted in chitin microcosms for 24 h at 30°C statically. Strains were distinguished by lacZ phenotype (using *lacZ*+ and Δ*lacZ* strain pairs) as previously described (16). Cultures were mixed in a 1:1 ratio and plated for quantitative culture on LB+Xgal. Competitive indices were calculated as previously described (16).

HGT assays between strain pairs were conducted with differentially marked strains where each contained an Ab^R^ marker at a distinct neutral locus (ΔVC1807::Erm^R^ or ΔVCA0692::Tm^R^). As indicated, strains were mixed in a 1:1 ratio in chitin microcosms for 48 h at 30°C statically (9). After 48 h, a portion of each co-culture was diluted in LB broth and outgrown for 2 hr prior to plating. Reactions were plated for quantitative culture on Tm+Erm plates to quantify transformants, as well as on Erm and Tm alone to quantify the abundance of each strain within the coculture. Transformation frequency is expressed as CFU of transformants (Erm^R^ + Tm^R^ double resistant) / (CFU of Erm^R^ + CFU of Tm^R^).

## ACKNOWLEDGEMENTS

The authors thank Dr. David Lazinski for scientific discussions. This work was supported by Fondecyt Iniciación en Investigación 11190049 (CAS-V), 11190158 (RCM-Q) and the National Institutes of Health R35GM128674 (ABD), and AI055058 (AC). Centro de Estudios Científicos (CECs) is funded by the Centers of Excellence Basal Financing Program of CONICYT PB-01.

## AUTHOR CONTRIBUTIONS

CAS-V, TND and RCM-Q performed experiments. CAS-V, RCM-Q, ABD and A.C. designed experiments and provided materials and strains. CAS-V, ABD and RCM-Q wrote the manuscript. All authors discussed the results and commented on the manuscript.

---

**Figure S1. Growth on insoluble chitin specifically induces K139 lytic replication in an estuarine environment. (A-C)** *V. cholerae* E7946 was grown in chitin microcosms with glucose (0.2%) and lactate (0.5%) added to growth reactions as indicated. Addition of these readily available carbon sources should diminish the reliance on chitin as a carbon source in these assays. After 24 h of static growth **A**, K139 PFU in culture supernatant and **B**, *V. cholerae* CFU were determined. **C**, K139 PFU generated per viable cell plotted as PFU/CFU using the data shown in **A** and **B**. (**D-F**) *V. cholerae* E7946 grown in M9 minimal medium with the sole carbon sources indicated: chitin (insoluble), glucose, lactate, the chitin monosaccharide N-acetyl glucosamine (GlcNAc), or soluble ß 1,4-linked GlcNAc oligosaccharides. After 24 h of static growth **D**, K139 PFU in culture supernatant and **E**, *V. cholerae* CFU were determined. **F**, K139 PFU generated per viable cell plotted as PFU/CFU using the data shown in **D** and **E**. Through these experiments, we found that the ratio of K139 viral particles per cell was highest during growth on insoluble chitin, thus, *V. cholerae* may specifically induce K139 production while utilizing insoluble chitin in microcosms. All data are from three independent experiments and plotted as the mean ± SD. Statistical comparisons were made by Student’s two-tailed t-test by comparing each condition to the control on chitin (*p<0.05, **p<0.01, ***p<0.001, ****p<0.0001).

