## Supplemental Table 1 for "Prophage-dependent neighbor predation fosters horizontal gene transfer by natural transformation"

**Table S1. Strains used in this study**

| Name / Strain# | Genotype | Reference |
|---|---|---|
| **E7946 WT / AC53** | Spontaneous SmR derivative of E7946, El Tor Biotype | Laboratory collection |
| **A1552** | Wildtype strain | Laboratory collection |
| **C6706** | Wildtype strain | Laboratory collection |
| **HC1037** | Wildtype strain | Laboratory collection |
| **ML111** | E7946 ΔlacZ::KanR | This study |
| **ML137** | E7946 ΔlacZ::KanR, ΔK139::CmR | This study |
| **TND2527** | E7946 ΔVCA0692::TmR, ΔlacZ::SpecR-PrecN-GFP | This study |
| **TND2525** | E7946 ΔVC1807::ErmR, ΔlacZ::SpecR-PrecN-GFP | This study |
| **TND2678** | E7946 ΔVCA0692::TmR, ΔlacZ::SpecR-PrecN-GFP , ΔK139::CmR | This study |
| **TND2543** | E7946 ΔVC1807::ErmR, ΔlacZ::SpecR-PrecN-GFP , ΔK139::CmR | This study |
| **TND2541** | E7946 ΔdprA::ZeoR, ΔVCA0692::TmR, ΔlacZ::SpecR-PrecN-GFP | This study |
| **TND2685** | E7946 ΔdprA::ZeoR, ΔVCA0692::TmR, ΔlacZ::SpecR-PrecN-GFP, ΔK139::CmR | This study |
| **TND2686** | E7946 ΔdprA::ZeoR, ΔVC1807::ErmR, ΔlacZ::SpecR-PrecN-GFP, ΔK139::CmR | This study |
