## Supplementary figures and images for "Prophage-dependent neighbor predation fosters horizontal gene transfer by natural transformation"

### Supplemental Figure 1

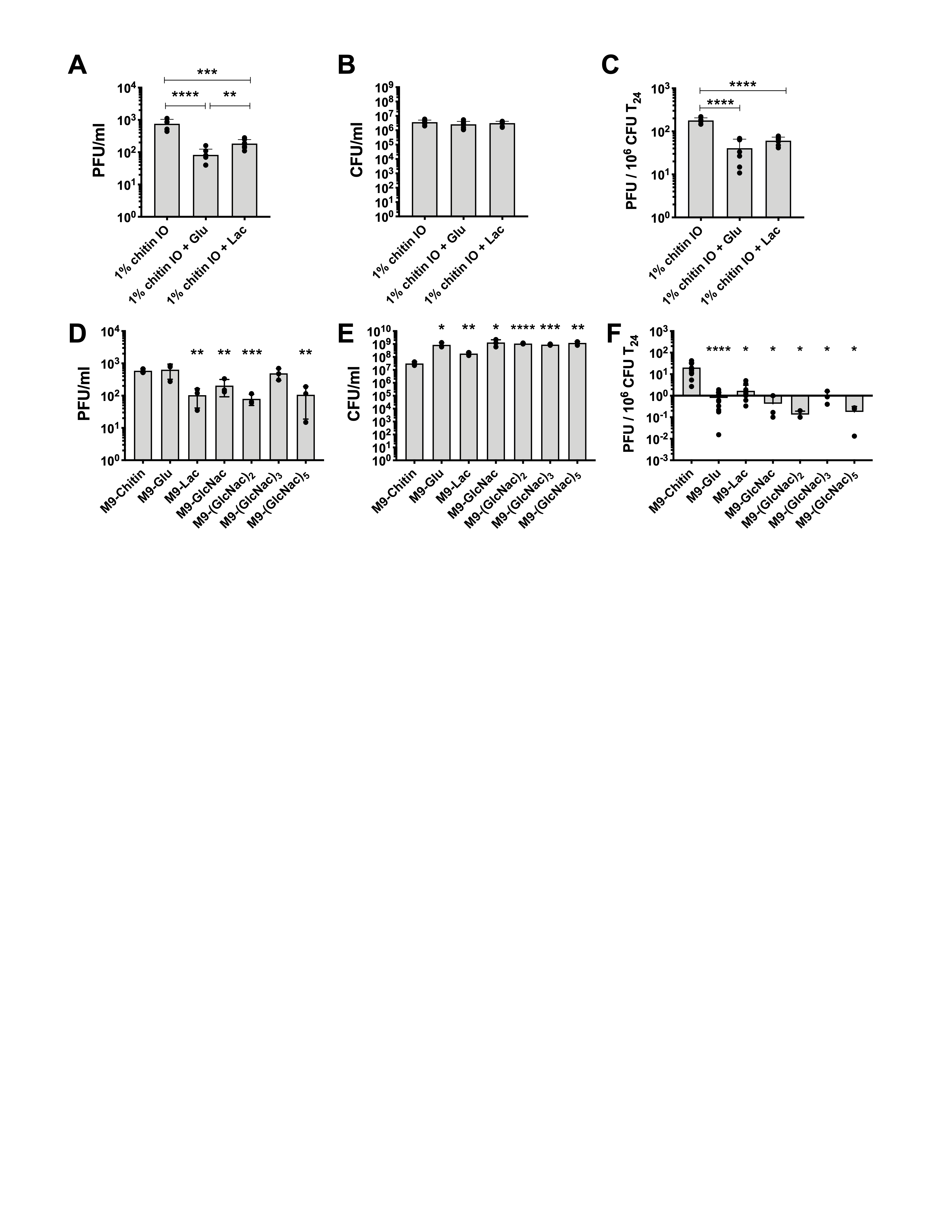
